# Long fragments achieve lower base quality in Illumina paired-end sequencing

**DOI:** 10.1101/397158

**Authors:** Ge Tan, Lennart Opitz, Ralph Schlapbach, Hubert Rehrauer

**Affiliations:** Functional Genomics Center Zurich, ETH Zurich/University of Zurich, Zurich, Switzerland

## Abstract

Illumina’s technology provides high quality reads of DNA fragments with error rates below 1/1000 per base. Runs typically generate a millions of reads where the vast majority of the reads has also an average error rate below 1/1000. However, some paired-end sequencing data show the presence of a subpopulation of reads where the second read has lower average qualities. We show that the fragment length is a major driver of increased error rates in the R2 reads. Fragments above 500 nt tend to yield lower base qualities and higher error rates than shorter fragments. We demonstrate the fragment length dependency of the R2 read qualities using publicly available Illumina data generated by various library protocols, in different labs and using different sequencer models. Our finding extends the understanding of the Illumina read quality and has implications on error models for Illumina reads. It also sheds a light on the importance of the fragmentation during library preparation and the resulting fragment length distribution.

## Introduction

Next generation sequencing (NGS) has become one of the most widely used technologies in biomedical research with a rapid development of new applications. Due to its efficiency and low financial cost, NGS has substantially improved many projects, including *de novo* genome assembly sequencing, resequencing, transcriptome profiling, sequence variants detection, epigenome profiling and chromatin interaction. Illumina platforms represent by far the most widely used technology for generating short reads. Current Illumina platforms produce millions to billions of reads in a single run with more than 75% of the reads reaching an average Phred quality score of Q30 which corresponds to a per base error rate of 1/1000. The high quality of the Illumina reads is among the reasons of Illumina’s success and can be verified readily by analyzing the wealth of Illumina data deposited in public repositories. An intriguing observation on the Illumina reads, however is that in paired-end sequencing the quality of the second read can be low and have a wide spread within a sample. It can be expected that the average quality of the second read is lower. However, the magnitude of the quality drop relative to the first read, can vary a lot between reads of the same library in the same run. As a consequence, a population of second reads with average quality well below Q30 may be observed even if basically all first reads have high quality. The size of this fraction of low quality reads can vary largely between runs, and even within the same run it may vary between multiplexed libraries of the same pool. Currently existing error models of Illumina reads do not provide an explanation of this phenomenon.

Previous studies have analyzed the characteristics of Illumina errors. It has been shown that sequencing errors are not completely random but exhibit systematic behaviors. The dominant error type of introduced errors are substitutions with very low rates of insertions and deletions and the read quality tends to be lower towards the end of the read (1). Certain preferences of substitutions errors have been also observed. Dohm et al. and Nakamura et al. reported wrong bases tend to be preceded by base G (1, 2). The frequency of the most frequent A to C conversion can be ten to eleven fold, over the least frequent conversion from C to G (1). Nakamura et al. also proposed that inverted repeats and GGC sequences patterns were the major causes of sequence-specific errors (2). Meacham et al. observed the systematic error of frequent sequence motif GGT due to substitution from GGG motif (3). A dependency of the quality on the base position in the read as well as strand-specific bias was also reported (4). Analyses of multiple datasets from several sequencers revealed the motifs that induce read errors are platform-dependent (5). However, none of the above findings explain the phenomenon of low read quality in subpopulations of the second read (R2 read) that we observed in some libraries. Although Soumitra et al. proposed a k-mer approach to filter out the error-containing reads at a high precision rate (6), a clear understanding of the source of errors from this paired-end sequencing is desired.

In this manuscript we demonstrate that fragment length impacts the error rates especially of the second read and does explain the observation of populations of second reads with low average quality. For this we determined fragment lengths by read alignment and stratified reads according to the fragment length. Subsequently, we computed and evaluated the Phred scores and base mismatch rates and their fragment length dependency. The analyses were done on sets of publicly available Illumina read data that we collected from the Illumina BaseSpace and the short read archive (SRA). We cover various sequencer models as well as library protocols. In the subsequent sections we report our results, discuss them, and describe out methods.

## Results

### Quality of the second read in paired-end sequencing

We surveyed publicly available paired-end sequencing data downloaded from Illumina BaseSpace and SRA that cover various library protocols and sequencer models (see Table 1). The identifier of the study is either the BaseSpace project name or the SRA project ID. In total, our dataset comprises 138 sequenced libraries from 27 different projects. In order to simplify read alignment we included only sequencing data from human or mouse samples.

**Table 1:** summary of investigated datasets

| Study | Protocol | Sequencer<br>Read length | Species | #Samples |
| --- | --- | --- | --- | --- |
| NovaSeq S1 Xp: Nextera DNA Flex (Replicates of NA12878) | WGS | NovaSeq6000 (2*151bp) | Homo sapiens | 1/4 |
| HiSeqX: Nextera DNA Flex (replicates of Coriell Trio Samples) | WGS | HiSeqX (2*151bp) | Homo sapiens | 1/24 |
| HiSeq 4000: TruSeq PCR-Free (NA12878) | WGS | HiSeq4000 (2*151bp) | Homo sapiens | 1/6 |
| HiSeq 2500: TruSeq PCR-Free DNA 2x251 (NA12878) | WGS | HiSeq2500 (2*251bp) | Homo sapiens | 1/2 |
| HiSeq X: TruSeq PCR-Free and Nano (UDI Replicates of NA12878 +/- Free Adapter Blocking Reagent) | WGS | HiSeqX (2*151bp) | Homo sapiens | 64/64 |
| NovaSeq S2: TruSeq PCR-Free 350 (Replicates of NA12878) | WGS | NovaSeq6000 (2*151bp) | Homo sapiens | 12/12 |
| HiSeq 4000: TruSeq PCR-Free and Nano (350bp to 550bp insert size) | WGS | HiSeq4000 (2*151bp) | Homo sapiens | 8/8 |
| HiSeq 4000: RNA-Seq 64-plex (MAQC HBRR and UHRR) | RNA-Seq | HiSeq4000 (2*76bp) | Homo sapiens | 1/64 |
| HiSeq 2500: TruSeq Stranded mRNA LT (SEQC: UHRR & Brain) | RNA-Seq | HiSeq2500 (2*75bp) | Homo sapiens | 1/4 |
| NovaSeq S2: TruSeq Exome (96 replicates of NA12878) | WES | NovaSeq6000 (2*76bp) | Homo sapiens | 1/96 |
| HiSeq 4000: TruSeq Exome (12plex replicates of NA12878) | WES | HiSeq4000 (2*76bp) | Homo sapiens | 1/48 |
| HiSeq 2500: TruSeq Rapid Exome (12 replicates of NA12878) | WES | HiSeq2500 (2*101bp) | Homo sapiens | 1/12 |
| SRP066955 | RNA-Seq | HiSeq4000 (2*50bp) | Mus musculus | 4/4 |
| SRP139105 | RNA-Seq | HiSeq4000 (2*151bp) | Mus musculus | 4/16 |
| SRP137053 | RNA-Seq | HiSeq4000 (2*126bp) | Mus musculus | 4/16 |
| SRP132952 | RNA-Seq | NovaSeq6000 (2*101bp) | Mus musculus | 4/12 |
| SRP124637 | RNA-Seq | NovaSeq6000 (2*150bp) | Mus musculus | 2/15 |
| SRP143964 | RNA-Seq | HiSeq2500 (2*150bp) | Mus musculus | 3/18 |
| SRP142426 | RNA-Seq | HiSeqX (2*150bp) | Mus musculus | 4/6 |
| SRP136620 | RNA-Seq | HiSeqX<br>(2*150bp) | Mus<br>musculus | 4/6 |
| SRP143392 | RNA-Seq | HiSeq2500<br>(2*101bp) | Mus<br>musculus | 4/9 |
| SRP124981 | WGS | HiSeq4000<br>(2*150bp) | Mus<br>musculus | 1/32 |
| SRP072373 | WGS | HiSeq4000<br>(2*100bp) | Mus<br>musculus | 1/1 |
| ERP016331 | WGS | HiSeq2500<br>(2*151bp) | Mus<br>musculus | 2/487 |
| SRP145593 | WGS | HiSeqX<br>(2*150bp) | Mus<br>musculus | 2/9 |
| SRP132517 | RNA-Seq | NextSeq500<br>(2*75bp) | Mus<br>musculus | 3/10 |
| SRP129935 | RNA-Seq | NextSeq500<br>(2*76bp) | Mus<br>musculus | 3/14 |

As a first step of quality evaluation, we show the fraction of reads that has low quality. Figure 1 shows that generally a larger fraction of the R2 reads has low quality. The two subplots show the result using two alternative measures of quality. Figure 1A is based on the Phred-scaled base quality values generated by Illumina’s base-calling software. It shows the fraction of reads that has a mean Phred score below 30. A Phred score of 30 corresponds to an error probability of 1/1000 bases. Figure 1B shows the fraction of reads that has an average mismatch rate above 1/100 bases after read alignment. The plots show the distribution of the fractions over all samples, and both plots confirm: the fraction of R2 reads below the quality threshold is higher and more variable than for R1 reads. The difference of the plots is that Figure 1A is derived from the predicted error as computed from the fluorescence signal generated by the sequencing by synthesis process, while Figure 1B is derived on the actual base mismatches of the aligned reads relative to the reference sequence. Obviously mismatches can be caused by sequencing errors as well as alignment errors or genotype differences of the sequenced samples relative to the reference sequence. We do not endeavor to discriminate the sources of the mismatches since our focus is on the relative difference of the quality of the R1 and R2 reads. It is important to note that only read type dependent differences in base quality can lead to the difference observed in Figure 1A and especially 1B. The other potential sources for mismatches do affect R1 and R2 reads in the same fashion and do not generate the differences that are observed in the fractions of low quality reads. In conclusion, it is safe to attribute the observed quality difference between the two read types to an elevated error rate of the R2 read. The remarkable characteristic is that the fraction of low quality R2 reads apparently varies between samples and leads to a broad density distribution while the density distribution of the R1 read is narrow. This points to an additional error-driving factor that may depend on the sample, library protocol or sequencer.

**Figure 1.**
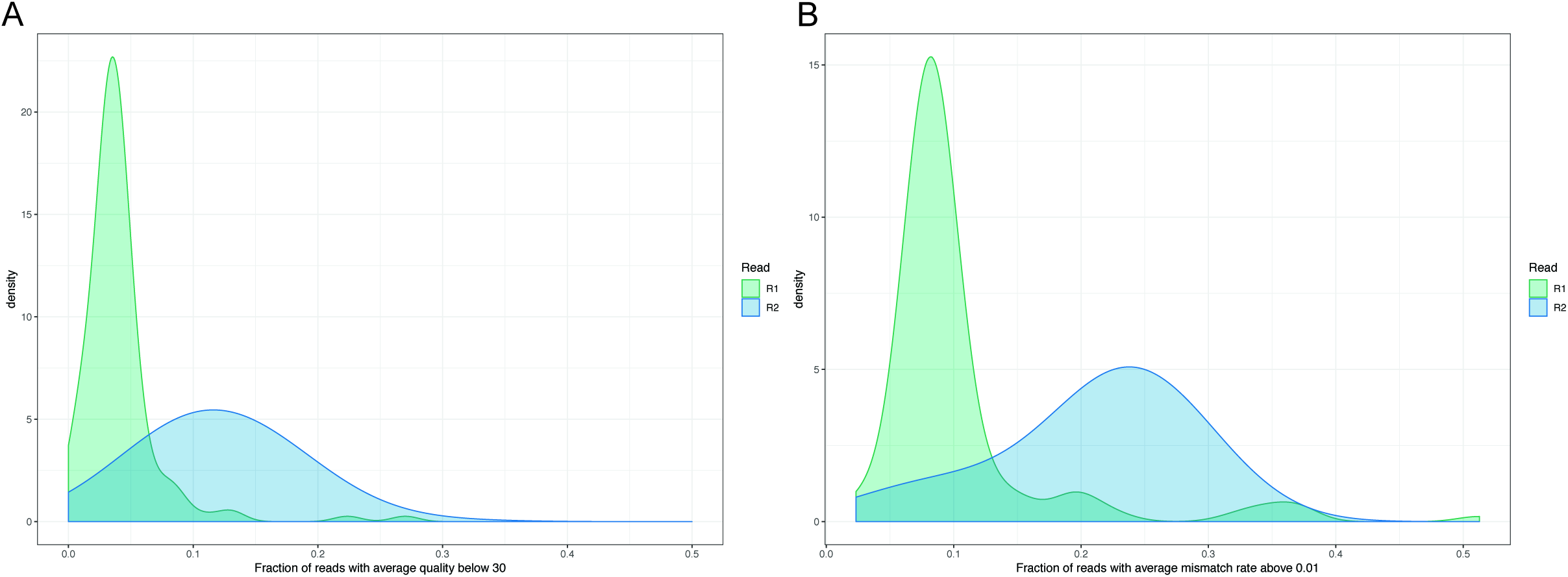
The read quality difference between R1 and R2 in Illumina paired-end sequencing. Higher fraction of R2 reads has low quality, compared with R1. A) Phred read quality based measurement. B) Mismatch rate based measurement.

### Fragment lengths and mismatch rates

When investigating what drives the magnitude of the quality drop of the R2 reads, we found that the fragment length distributions of the libraries have a major impact. Specifically, the fraction of low quality R2 reads is related to the fraction of long fragments in the library. As Figure 2 shows, the amount of additional base mismatches in the R2 read relative to the R1 read is correlated with the amount of fragments that have a length above 500 bp. Figure 2A shows that if the library has basically no long fragments then the R2 read has not a systematically higher mismatch rate than the R1 read. For samples where more than 10% of the fragments are longer than 500 bp, the R2 reads have more base mismatches than the R1 reads. Figure 2B compares additionally the absolute fractions of low quality reads in R1 and R2 in a scatter plot and indicates for each sample the fraction of long fragments by the coloring of the plotting symbols. The plots in Figure 2 also show that this characteristic is observed for all library types and all sequencer models analyzed. Notably, the data analyzed includes two experiments with samples where the library preparation and sequencing was identical and the targeted fragment length (350, 450 and 550 bp) is the only parameter that has been varied. The results for these samples is shown as connected lines in Figure 2. Figure 2B shows that for those samples the fraction of low quality R2 reads strictly increases with the fraction of long fragments. The trend observed in those samples corroborates that the low R2 qualities are driven by fragment length and not by the tissue source, library type, or sequencer model. Figure 2B shows also that for four RNA-seq experiments the R1 read has a higher fraction of low quality reads than the R2 reads which comes as an unexpected result. It is known that Illumina’s RNA-seq protocols lead to lower quality bases at the read start (5’-end of the fragments) (7). A closer inspection revealed that in those four samples this 5’-end low quality effect is very strong, actually stronger than the R2 quality drop (Supplementary Figure 1). Since the fragments are longer than the read length the low quality 5’-end of the cDNA only affects the R1 read but not the R2 read.

**Figure 2.**
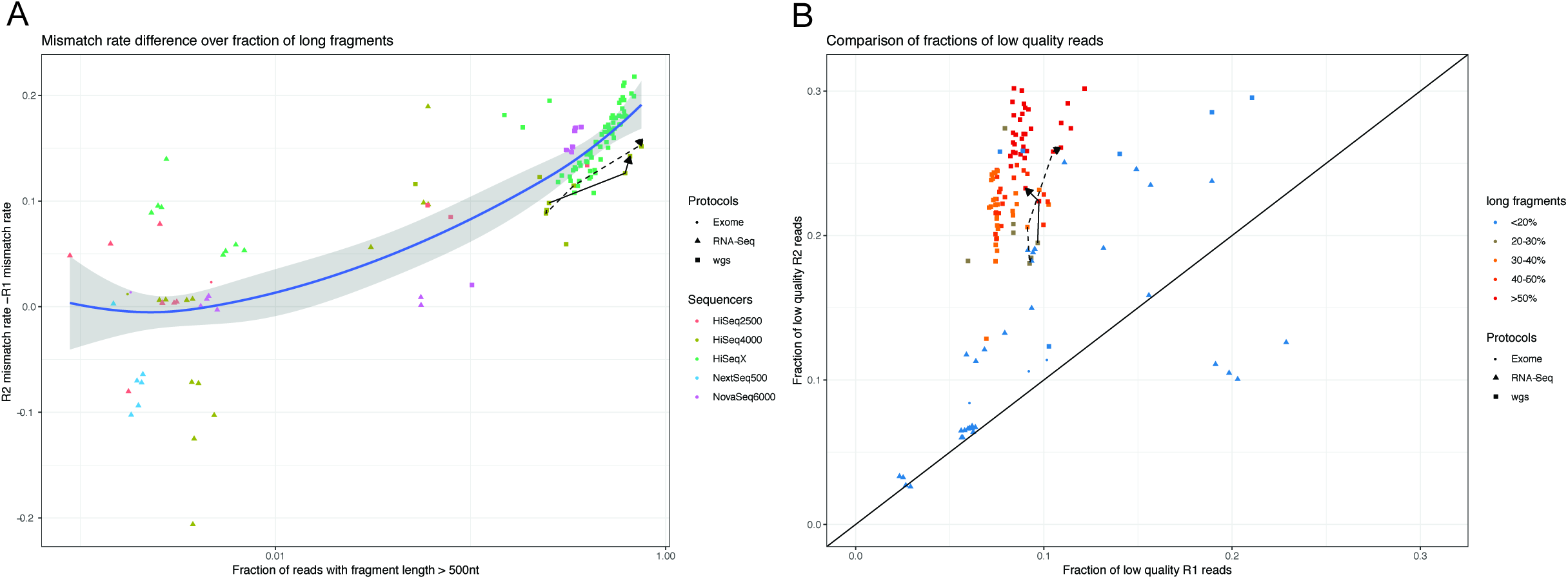
The impact of fragment length on the drop of R2 read quality in paired-end sequencing across multiple sequencers and various protocols from BaseSpace datasets. Two experiments with varied insert size are connected with lines. A) For almost every dataset, the fraction of low quality R2 reads is higher than the fraction of low quality R1 reads. With increasing fraction of long fragments (>500 nt), particularly, there are more low quality R2 reads than R1 reads, regardless of the protocols. B) The difference of R2 mismatch rate and R2 mismatch rate gets higher along with increasing fraction of long fragments (>500 nt), regardless of the sequencers and protocols. The mismatch rate is calculated as the mean of the average read qualities.

In order to assess the details of the dependency of base and read quality on the fragment length, we stratified the reads according to fragment length and analyzed the strata separately. Figure 3 shows the fraction of low quality reads for different fragment length strata. The plot shows that for the R1 read, the fraction of low quality reads has no strong systematic dependency on the fragment length. Reads are reliable for fragments ranging from 200 to 1000 bp. In contrast to this, the fraction of low quality reads increases clearly for fragments in the range of 500 to 600 bp and above. Again the plot shows that this trend is observable for different sequencer models and library types.

**Figure 3.**
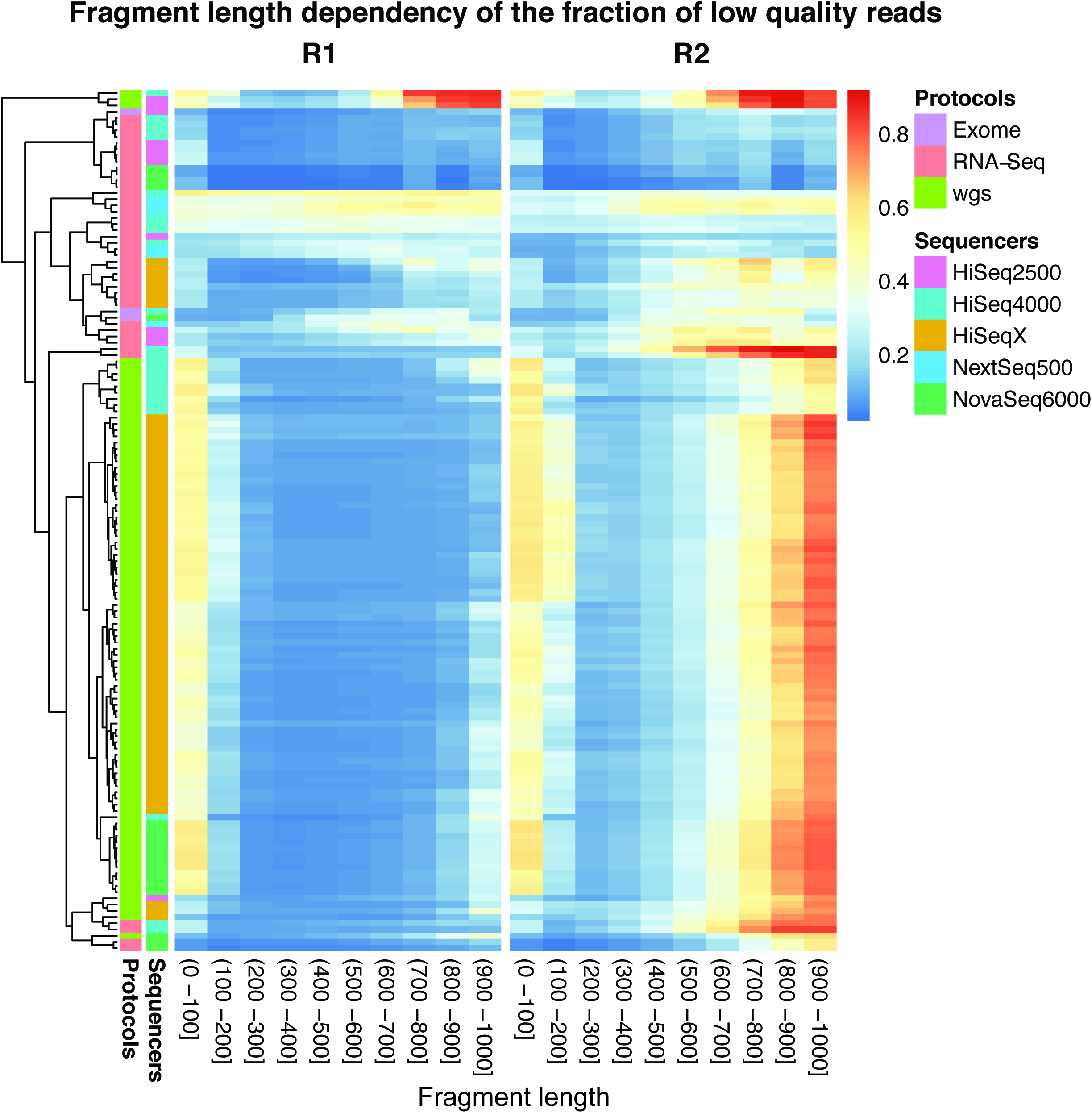
Fraction of low quality reads from each fragment length strata across various sequencers and protocols. Compared with R1, the fraction of low quality reads in R2 has stronger dependency on the fragment length. The read quality drops considerably for fragments longer than 500 nt.

### Dependency of base quality on base position

We additionally verified the positions of the low quality bases and characterized the distribution of low quality positions. It is known that Illumina’s sequencing by synthesis leads to a quality decrease as more and more bases are sequenced. This is caused by the fact that phasing errors occur within a cluster and accumulate (1). At every sequencing step, individual molecules of a cluster may fail to get a newly synthesized base. If the number of erroneously synthesized molecules in a cluster increases the sequencer detects a mixed signal which can lead to base calling errors. In Figure 4, we plot the average mismatch rates per base position for short fragments (200 - 400 bp) and long fragments (600 - 1000 bp). Figure 4A and 4B show the mismatch rates for the R1 and R2 read separately. Figure 4A confirms the known behavior of decreasing quality towards the end but also shows that the R1 quality of long fragments is very close to the quality of short fragments. Figure 4B, however, shows that the R2 follows a different pattern. Already the early bases in the long fragments have a mismatch rate that is clearly above the mismatch rate observed for short fragments. Interestingly the length-dependent increase in the mismatch rate for the long fragments is also more pronounced than for the short fragments.

**Figure 4.**
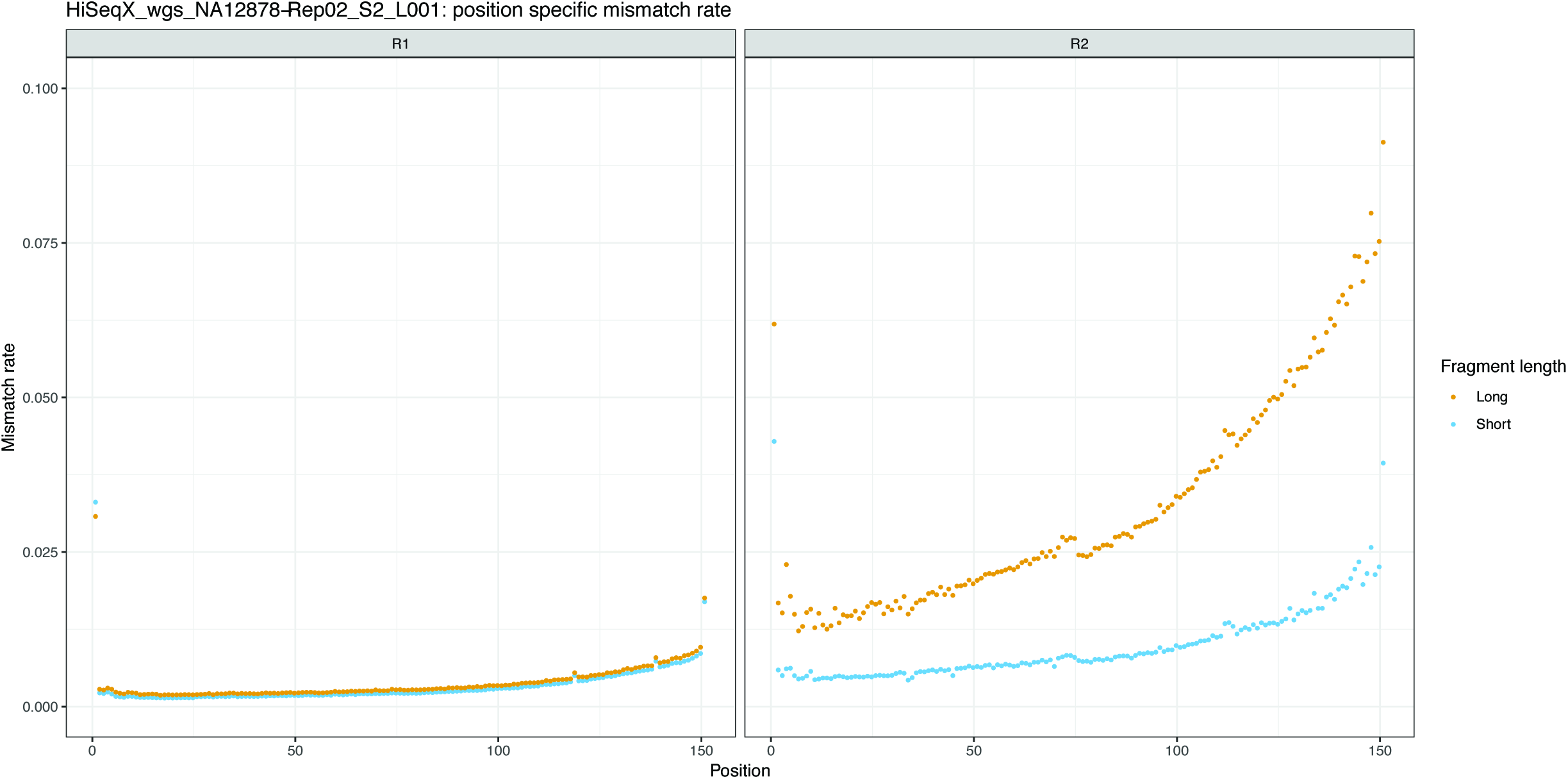
Position specific averaged mismatch rate for short and long fragments in R1 and R2. One example of sample NA12878 from HiSeqX platform with WGS protocol is shown here. In R1, the short and long fragments exhibit similar position specific mismatch rate, while in R2, the long fragments lead to much higher mismatch rate than short fragments.

## Discussion

This is the first time that the impact of the fragment length on the base qualities of Illumina reads has been investigated and quantified. Previous research has looked at various other factors like sequence context, base position in read, *etc.*, but none of these does explain the widely observed lower quality in the R2 read in sub-populations of reads from the same sample on the same sequencing run. Our analysis identifies the fragment length as the major cause for this higher risk of low quality of an Illumina R2 read relative to the corresponding R1 read of the same fragment. We find that if libraries have higher fractions of long fragments, then the reported Phred scores tend to be lower and the mismatch rates in paired-end alignment tend to be higher. Overall our analysis confirms that the Phred qualities are good representatives of the actual base quality. We find also that sequencing quality is optimal if libraries are prepared according to Illumina’s specifications with short fragment lengths like 250 bp for RNA-seq and 350 bp for DNA-seq. We can also confirm that the presence of low quality reads from long fragments does not affect the high quality of reads achieved for the shorter fragments in the same library that are in the range of 200 to 400 bp. Further, we observe rather lower and highly variable quality for very short fragments in the range of 0 to 100 bp. However, here our evaluation may be compromised by failed trimming of partial adapters at the 3’-end. Additionally, usually libraries have only very few fragments in that size range.

Our conclusions do not rely on an analysis of the Phred-scale qualities computed by the Illumina software but rather on the base mismatches after paired-end alignment. By that our results are independent of proprietary software and we can guarantee that our computations do not consider the sequencer model in any form. We only make statements on the differences of these mismatch rates between short and long fragments and the differences between the first and the second read. With this approach, mismatches in the alignment that are caused by genotype differences and misalignments do cancel out. This means, by focusing on the differences of these mismatch rates, we eliminate potential genotype and alignment effects. Our study shows indeed that the mismatch rate varies between samples in different studies, but the observed increase of the mismatch rate in the R2 read of long fragments is stably observed in all studies. Since our study suggests that the optimal quality of Illumina reads is achieved if the input library has only short fragments, the protocols that intrinsically generate short fragments are less affected. These protocols include, e.g., whole exome sequencing protocols where the enrichment of exonic fragments uses sheard input DNA in the range of 150 - 200 nt. For all samples investigated, we observed a high mapping rate. This suggests that in applications where aligned reads are finally counted and only the read counts are used, the reads from the long fragments are still used for purpose. The fragment length effect on quality limits the choices of fragment lengths in *de novo* sequencing projects. For those, longer fragments would help the assembly but are not advisable due to the increased error rate. Especially, due to the fact that one would need a paired-end alignment in order to determine the fragment length but that would not be available in a *de novo* setting.

Based on our results, a fragment length filter as a preprocessing step of variant calling might be useful in situations where low frequency mutations are searched in heterogeneous cell populations or where variants are called from low coverage sequencing data. The results give also indications that the modelling of Illumina errors (for a review of read error modelling see (8)) can be improved by including fragment length as a factor. Providing explanations on the actual origin of the quality drop for long fragments is beyond the scope of this study. However, the fragment-length independent high quality of the R1 reads suggests that none of the steps until sequencing the R1 read is affected. This does imply that cluster amplification works equally well for short and long fragments.

## Materials and Methods

### Data processing and mapping

Reads were preprocessed by Trimmomatic (9) (ILLUMINACLIP:adapters.fa:1:20:7:5:true MINLEN:20) in order to remove adapters from the 3’-end. Subsequently reads were aligned with Bowtie2 (10) (–no-unal -D 20 -R 3 -N 1 -L 20 -i S,1,0.50 –maxins 1000 –minins 0). We chose to be error-tolerant in order to map as many reads as possible. The sensitive options will also align the reads with many errors. Being stringent would have implied that low quality reads were discarded which would have biased our result. The parameter choices will report concordant paired-end alignments only if the fragment length is 1000 bp or below. We found that this does not represent a limitations since in all libraries the number of fragments with larger size were very low. In fact so low that the we considered them to few in order to draw conclusions. SRA and BaseSpace were aligned against mouse (GRCm38.p5) and human (GRCh38.p10) references, respectively. The whole-genome sequencing reads were aligned to reference genome, while RNA-Seq reads were aligned against the corresponding set of transcripts downloaded from Ensembl. We chose transcriptome alignment in order to be able to use the same aligner as for the DNA samples and still have a good estimate of fragment length. This choice implies that reads that come from splice variants not represented in the transcriptome will be omitted. In order to minimize errors caused by alignment errors, we look only at the primary alignments. And since we inspect the dependency on fragment length we only look at properly paired reads. For all others we would not know the fragment lengths. Aligned BAM files were processed with samtools (11) to produce MD tags and identify the bases with mismatches. All computations and plots were computed with R scripts.

## Acknowledgements

We are thankful to the sequencing and the genome informatics team of the FGCZ for the helpful discussions and the great working atmosphere.

